# Familial Alzheimer’s disease mutation undermines axonal transport by enhancing dynactin recruitment to the APP motor assemblies

**DOI:** 10.1101/2022.11.17.516829

**Authors:** Monica Feole, Gorazd B. Stokin

## Abstract

Experiments in flies, mice and humans suggest a significant role of impaired axonal transport in the pathogenesis of Alzheimer’s disease (AD), however, the underlying mechanisms remain unknown^1,2^. We report that the Swedish familial AD (FAD) mutation perturbs fast anterograde axonal transport of the amyloid precursor protein (APP) by altering directionality of its movement. APP thus spends more time in retrograde movement and accumulates in the soma. We found that the Swedish mutation enhances recruitment of dynactin 1 to the APP transport assemblies. Given that dynactin 1 activates the retrograde motor dynein^3^, this hampers physiological anterograde axonal transport of APP. We last show that the Swedish mutation perturbs also the axonal transport of early endosomes, which rely on the same molecular motors as APP. Our findings reveal extensive impairment of the axonal transport pathways by a FAD mutation, which reflects dysregulation of the cargo motor assemblies.

## INTRODUCTION

The APP is a type I integral membrane protein active at synapses^4^ and best known for its role in the amyloid pathology and pathogenesis of AD^5^. In fact, autosomal dominant mutations in APP have long been identified to segregate with kindreds afflicted by AD^6^. Although exceptionally rare, these familial AD (FAD) mutations play an invaluable role in elucidating mechanisms underlying the pathogenesis of AD. For example, the APP KM670/671NL Swedish double mutation promotes β-cleavage of APP at the N-terminus of its amyloid-β peptide (Aβ) sequence^7^. This cleavage enhances formation of β-cleaved APP C-terminal fragments (β-CTFs), which are subject to γ-cleavage at the C-terminus of the Aβ sequence and release excess of Aβ^8,9^. Other FAD mutations, such as the APP V717I London mutation, promote γ rather than β-cleavage and also release excess of Aβ^10,11^. Aberrant Aβ production spearheads the amyloid cascade hypothesis, which postulates that Aβ ignite and drive AD pathogenesis^12^. FAD mutations, however, increase also β-CTFs levels and enlarge early endosomes, which suggests that mechanisms of perturbed intracellular sorting and degradation are likewise at play in the pathogenesis of AD^13,14^.

In axons, APP undergoes fast anterograde transport^15^ and proteolytic cleavage into β-CTFs and Aβ^16^. Although interactions between the components of the APP motor assemblies remain to be further elucidated^17,18^, a number of studies reports that APP vesicles move within the axons by highly processive molecular motors, kinesin-1 in the anterograde and dynein-dynactin complex in the retrograde direction^19-21^. Studies in animal models and patients afflicted by AD point to a role of axonal transport and pathology in the pathogenesis of AD and suggest that FAD mutations perturb axonal transport^22-26^. These studies gain further support from cell culture experiments, which demonstrate that FAD mutations reduce proportion of anterogradely transported APP^27^. However, changes in transport behaviour of the cargoes and in the processivity of molecular motors that would instruct about mechanisms underlying putative transport disruption by FAD mutations remain unknown. We here rigorously characterize the effects of FAD mutations on axonal transport and provide an insight into the mechanisms by which FAD mutations impair axonal transport.

## RESULTS

### FAD mutations impair anterograde axonal transport of APP in human neurons

To investigate the impact of FAD mutations on APP transport, we first recreated previously reported experimental settings using human neurons^27,28^. Neurons were transfected with either wildtype APP (APP_wt_) or APP harbouring Swedish (APP_swe_) or London (APP_lon_) mutations, linked to Green Fluorescent Protein (GFP). Two days following transfection, movies of APP transport were acquired from distal neuronal projections (Movies S1-3). APP_swe_ and APP_lon_ exhibited significant reduction in the proportion of anterogradely transported particles compared to APP_wt_ (Fig. S1a-c).

To delve into mechanisms underlying the observed impairment in transport by FAD mutations, we focused on the Swedish mutation and first established its overall effect on the axonal transport of APP. Neurons were grown for 40 DIV in ibidi multichannel devices and then transduced with either APP_wt_, linked to GFP, or APP_swe_ linked to Red Fluorescent Protein (tRFP) (Fig. 1a). Ten days following transduction, movies of APP transport were acquired from distal neuronal projections. Neuronal cultures were then stained for axonal (pNFH) and dendritic markers (MAP2) (Fig. 1b). Using position retrieval, we selected for further analysis only those movies of APP transport in the distal projections that stained with the axonal marker (Fig. 1c, Movies S4 and 5). Semi-automated tracking algorithm was then employed to measure net movement of APP. Measurements showed significant reduction in the proportion of anterogradely transported APP_swe_ compared with the APP_wt_ particles with no differences in the number of tracks (Fig. 1d-f, Movies S6 and 7). This reduction was accompanied by significant increase in the stationary APP_swe_ particles. APP_swe_ also showed significant decrease in anterograde, but not retrograde, average velocity of axonal transport compared with APP_wt_ (Fig.1g).

**Fig.1.**
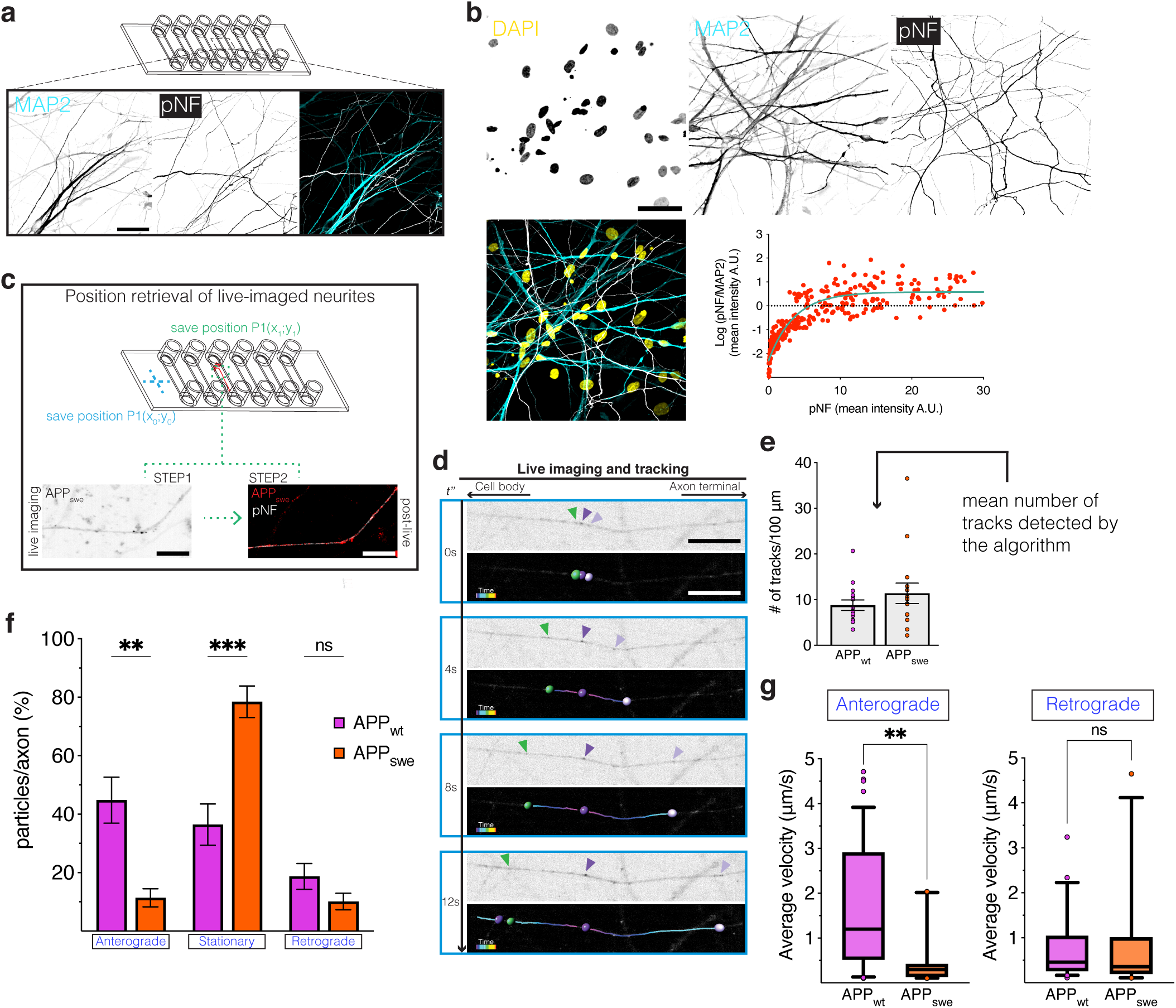
APP axonal transport is impaired by FAD mutations **a.** Representative image of pNFH and MAP2 immunostainings of human neurons cultured in ibidi multichannel (scale bar = 50µm). **b.** Mean intensities of pNFH/MAP2 ratios in neurons (n=303 neurites from 3 biological replicates). **c.** Schematic of the position retrieval approach used in ibidi multichannel; lower panel shows pictures of the same axon during live- (left) and post-imaging (right) stages; overlap between pNFH (+) and APP_swe_-tRFP (+) intensities (on the right, scale bar = 20 µm). **d.** Time frames of APP particles moving toward the axon terminal (light purple), cell body (green) or stationary (dark purple), each frame consists of live (above) and processed image (below) via semi-automated tracking (scale bar=20µm). **e.** Number of tracks/100µm quantified using semi-automated tracking for both APP_wt_ and APP_swe_; n=15 axons from 3 biological replicates. **f.** Proportions of APP_wt_ and APP_swe_ moving in anterograde direction, retrograde direction or stationary; n=15 axons from 3 biological replicates. **g.** Average velocities of anterogradely or retrogradely moving APP_wt_ and APP_swe_; n>10 particles per condition from 3 biological replicates. Data are mean±s.e.m. **(e, f)** or 10-90 percentile’s box-and-whiskers **(g)** (**p<0.01, ***p<0.001). Statistical comparisons were performed using unpaired *t*-test **(e)**, 2-way ANOVA followed by Šídák’s multiple comparisons **(f)** and Mann-Whitney test **(g)**.

To confirm that the observed transport behaviour of APP_swe_ is indeed of exclusively axonal nature, we validated our results also in microfluidic chambers (Fig. S1d, e). In conclusion, by employing several experimental paradigms designed to examine exclusively axonal transport, we demonstrate that the Swedish mutation impairs axonal transport of APP.

### Swedish mutation perturbs processivity of APP motor assemblies

To establish the precise behavioral changes in APP transport caused by the Swedish mutation, we performed an in-depth analysis of the movement of APP particles in axons (Fig. 2a). Segmental analysis showed a significant increase in the percentage of APP_swe_ particles in retrograde motion accompanied by their decrease in anterograde motion compared to APP_wt_, while pausing time remained unchanged (Fig. 2b). Intriguingly, only distances of APP_swe_ particles moving toward the presynaptic terminals, but not toward cell bodies, were shorter compared with APP_wt_ (Fig. 2c, Fig. S2a).

**Fig.2.**
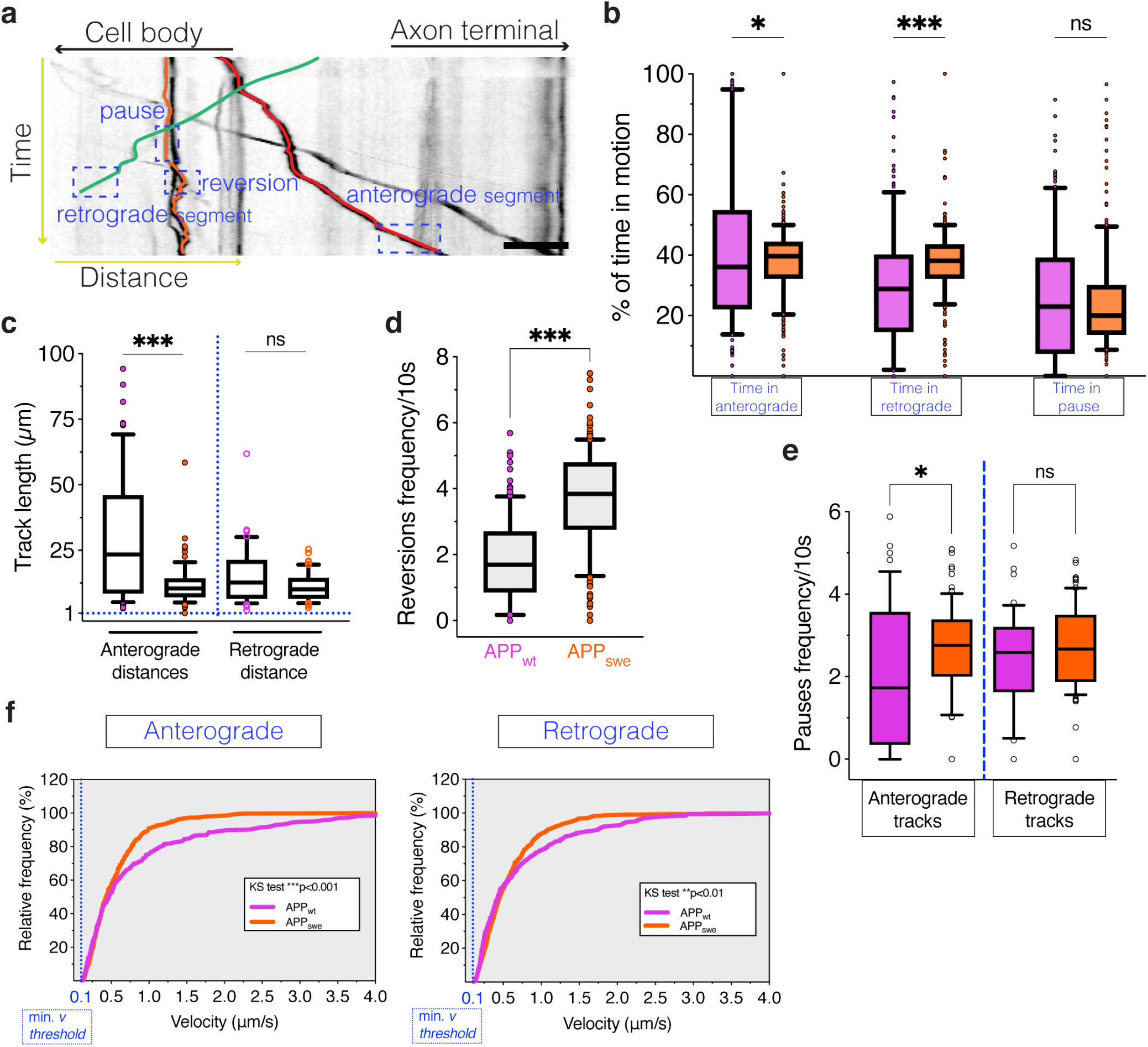
Swedish FAD mutation increases reversions and pauses frequencies **a.** Kymograph illustrating 2D trajectories (red=anterograde, green=retrograde, orange=stationary); dashed boxes showing the main axonal transport parameters based on the segmental analysis (scale bar=10µm). **b.** Real-time movement of APP_wt_ and APP_swe_ in anterograde and retrograde direction or pausing; n>150 particles from 3 biological replicates. **c.** Distances reached by APP_wt_ and APP_swe_ in anterograde or retrograde direction; n>45 particles from 3 biological replicates. **d.** Reversion frequencies/10s of APP_wt_ and APP_swe_; n>150 particles from 3 biological replicates. **e.** Pauses frequencies analysed for anterograde and retrograde tracks of APP_wt_ and APP_swe_; n>45 particles from 3 biological replicates. **f.** Cumulative frequency distribution of segmental velocities of APP_wt_ and APP_swe_ in anterograde and retrograde directions; n>400 segments from 3 biological replicates. 10-90 percentile’s box-and-whiskers **(b-e)** or cumulative frequency distributions (**f**, *p<0.05, **p<0.01, ***p<0.001). Statistical comparisons were performed using 2-way ANOVA followed by Šídák’s multiple comparisons test **(b)**, Mann-Whitney test **(c-e)**, and Kolmogorov-Smirnov test **(f)**.

In light of the overall reduced anterograde transport and distances and to better characterise the increased time spent in retrograde motion, we next investigated directionality of APP transport. Frequencies of reversions between anterograde and retrograde movement direction were significantly increased in APP_swe_ particles overall as well as in anterograde and retrograde tracks compared with APP_wt_ (Fig. 2d, S2b). On the other hand, pauses frequency, defined as the number of pauses which particles experience during their journey along the axon, showed significant increase only in the anterogradely transported APP_swe_ particles with no overall changes between APP_wt_ and APP_swe_ (Fig. 2e, S2c).

Increased pauses frequency in the anterogradely transported APP_swe_ particles, together with reduced anterograde transport, shorter track lengths and enhanced reversions, led us to next examine segmental velocities. Cumulative distribution analysis showed significantly reduced anterograde as well as retrograde segmental velocities of APP_swe_ compared with APP_wt_ particles (Fig. 2f). Reduction in segmental velocities was greater in anterograde (app. 0.3 µm/s) than in the retrograde direction (app. 0.15 µm/s, Fig. S2d). The above described behaviour of APP_swe_, characterized by bi-directionality of the transport due to accentuated retrograde movement, is indicative of the Swedish mutation perturbing processivity of the APP motor assemblies.

### Enhanced localization of APP carrying Swedish mutation in the soma

The observed predominantly anterograde impairment of axonal APP transport triggered by the Swedish mutation raises the question of whether APP accumulates in the cell bodies. To address this question, 40 DIV human neurons were fixed and stained for either GFP or tRFP, which mark APP_wt_ and APP_swe_, respectively, as well as with Tau to compare intensities between cell bodies and neurites (Fig.3a, b). APP_swe_ demonstrated significantly increased ratio of cell body *versus* neurite intensities compared with APP_wt_ (Fig. 3c). In contrast, there were no changes in the cell body to neurite Tau ratios in both APP_wt_-GFP and APP_swe_-tRFP transduced cell cultures (Fig. 3d). Our findings indicate that impaired anterograde axonal transport of APP by the Swedish mutation causes accumulation of APP in the cell bodies.

**Fig.3.**
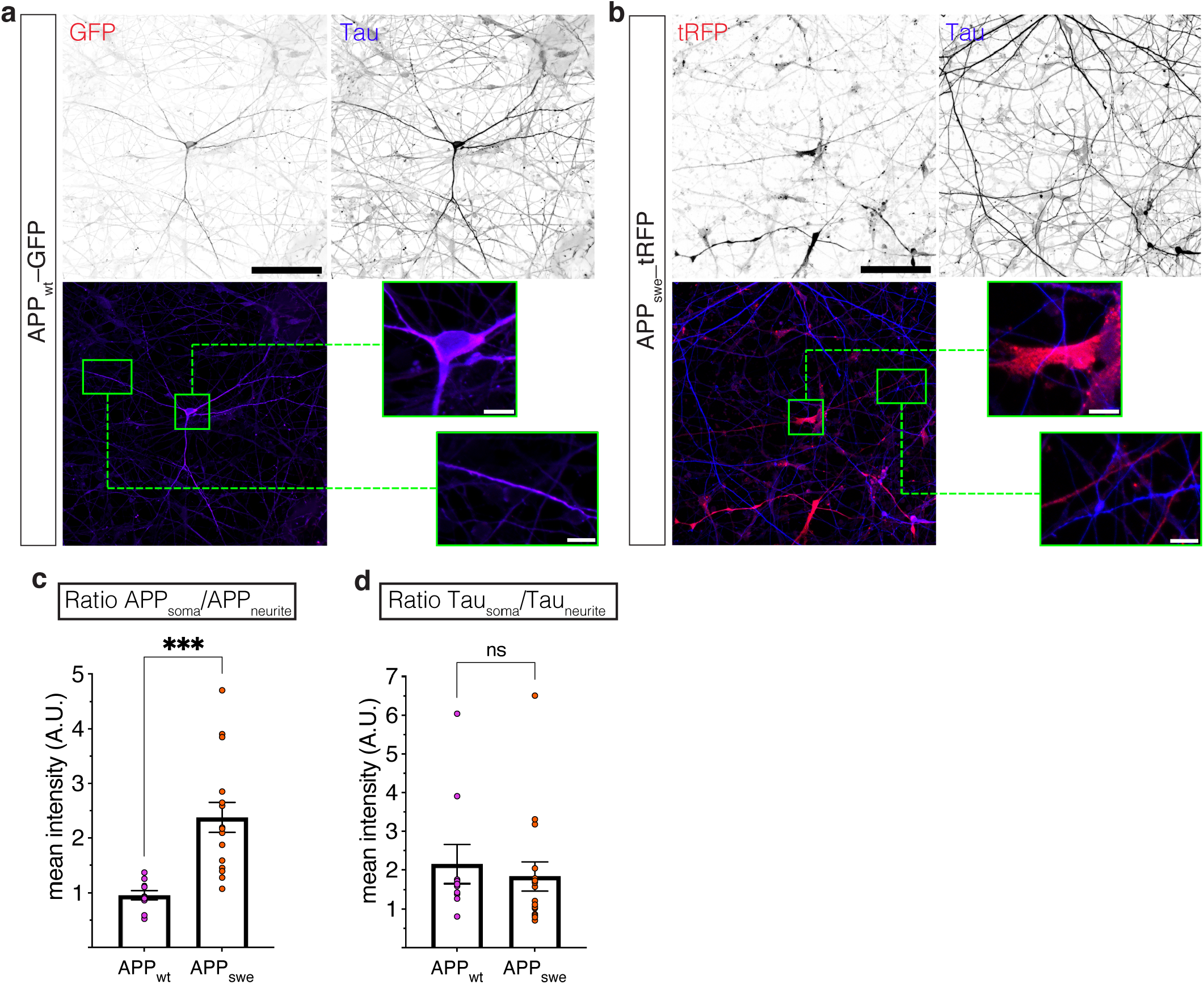
Swedish mutation perturbs APP distribution along neuronal projections Representative images of APP_wt_ **(a)** or APP_swe_ **(b)** transduced mature human neurons stained against either GFP or tRFP and Tau (scale bar=100 µm); zoom-in images for both cell bodies and distal projections (scale bar=1µm). **c.** Mean intensity ratios of APP_wt_ or APP_swe_ signals in soma/projection; n≥10 pictures from 3 biological replicates. **d.** Mean intensity ratios of Tau signals in soma/projection in the same ROIs as in **(c)**; n≥10 pictures from 3 biological replicates. Mean±s.e.m. (***p<0.001). Statistical comparisons were performed using unpaired *t*-test.

### Swedish mutation enhances recruitment of dynactin 1 to the APP motor assemblies

Evidence of perturbed processivity of APP motor assemblies by the Swedish mutation is suggestive of impaired coordination of APP movement by the molecular motors. To test for changes in the interaction between the components of the APP motor assemblies, we transduced 20 DIV SH-SY5Y cells with either APP_wt_ or APP_swe_. Cell lysates were next immunoprecipitated (IP-ed) with either GFP or tRFP, which tag APP_wt_ and APP_swe_, respectively, and separated on SDS-PAGE. The blots were then probed for detection of APP, kinesin light chain 1 (KLC1) and dynactin 1 (DCTN1) (Fig. 4a, S3a). The ratio of intensities between APP and DCTN1, but not KLC1, significantly increased in lysates from cells expressing APP_swe_-tRFP compared with APP_wt_-GFP (Fig.4b). Considering the same membranes re-blotted for either GFP or tRFP recognized exclusively over-expressed APP, while IP of non-transduced (NT) cultures showed no binding with antibodies against GFP or tRFP, our results demonstrate that the increased ratio of intensities between APP and DCTN1 is the consequence of the Swedish mutation (Fig.S3b and c).

**Fig.4.**
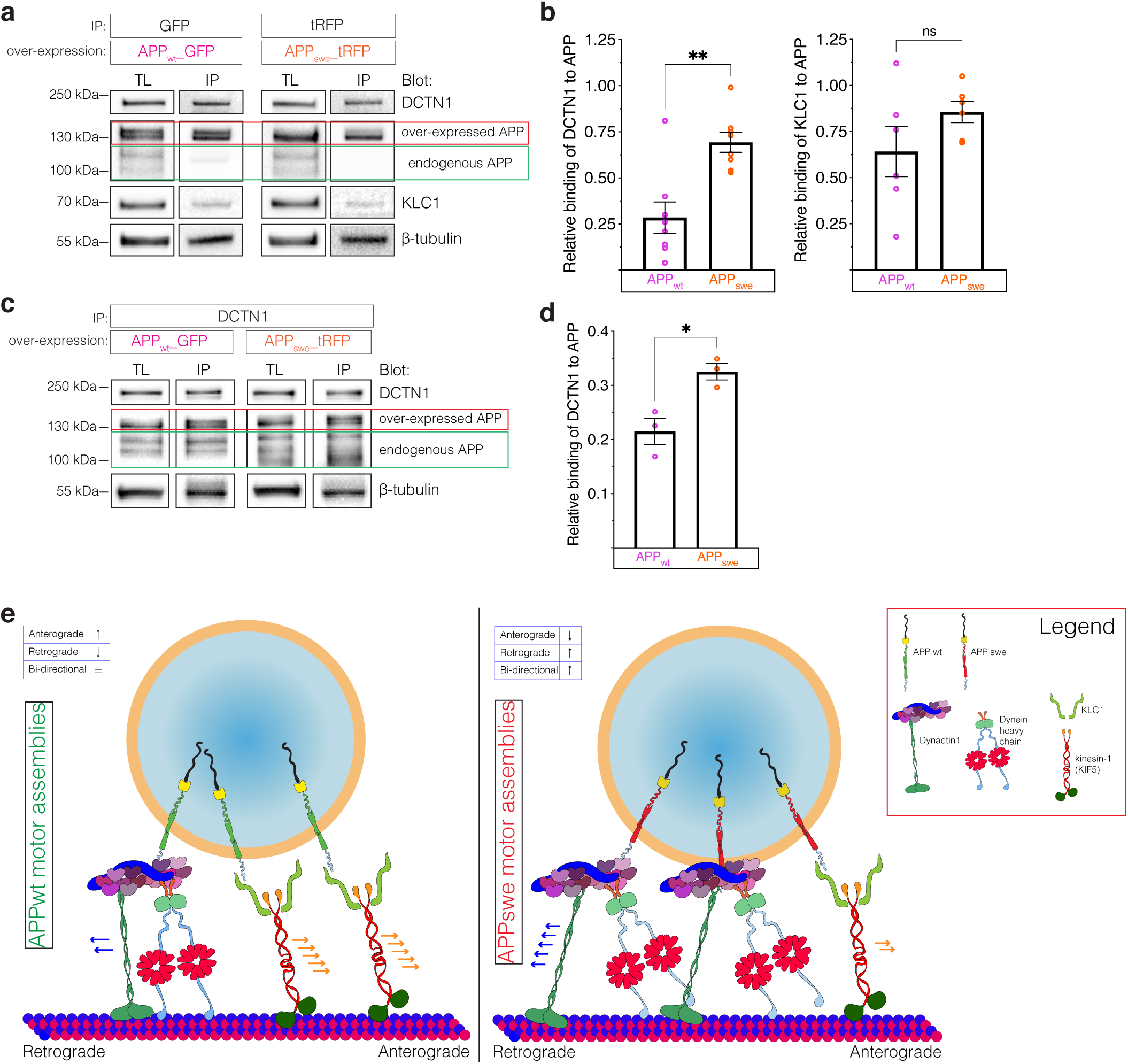
Enhanced recruitment of Dynactin1 to the APP motor assemblies **a.** Representative images of GFP and tRFP IPs of APP_wt_ or APP_swe_ lysates with membranes blotted for APP, DCTN1, KLC1 and βIII-tubulin (loading control). **b.** Intensity ratios of DCTN1 or KLC1 levels relative to either APP_wt_ or APP_swe_; n=8 (DCTN1 co-IPs) and n=6 (KLC1 co-IPs). **c.** Representative images of DCTN1 IP of APP_wt_ or APP_swe_ lysates with membranes blotted for APP and βIII-tubulin (loading control). **d.** Intensity ratios of APP levels relative to DCTN1; n=3 DCTN1 Ips. **e.** A model showing the consequences of enhanced DCTN1 recruitment to the APP motor assemblies. Mean±s.e.m. (*p<0.05, **p<0.01). Statistical comparisons were performed using unpaired *t*-test.

To test further for enhanced recruitment of DCTN1 to APP motor assemblies, we IP-ed SH-SY5Y cell lysates with DCTN1 and probed the blots with antibody against APP (Fig.4c, S3d). We found significantly increased ratio of intensities between DCTN1 and APP in cells transduced with APP_swe_ compared with APP_wt_ (Fig.4d). Collectively, these experiments show that Swedish mutation enhances recruitment of DCTN1 to the APP vesicles. Considering DCTN1 plays a role in a activating dynein^3^ our findings suggest increased activation and processivity of the dynein-dynactin complex. (Fig.4e).

### Swedish mutation promotes anterograde axonal transport of early endosomes

Considering molecular motors multitask in transporting several cargoes along the axons, we asked whether enhanced recruitment of DCTN1 to APP assemblies perturbs axonal transport of other cargoes driven by the same motors. To this end, we selected to study early endosomes since they are transported by the same molecular motors as APP and participate in the pathophysiology of AD^29^. To first reproduce the previously reported phenotype of increased size of Rab5 positive (Rab5+) endosomes in human induced pluripotent cell lines carrying FAD mutations^13,30^, we transduced human neurons with APP_swe_ and stained for Rab5 and tRFP to acquire high-resolution images of neurons expressing APP_swe_ (Fig. 5a). Neurons expressing APP_swe_ showed a significant increase in size of the area occupied by the Rab5+ particles with reduced frequency of endosomes with areas ≤ 0.5 µm^2^ and increased frequency of endosomes with areas ≥ 0.5 µm^2^ compared with NT cultures (Fig. S4a).

**Fig.5.**
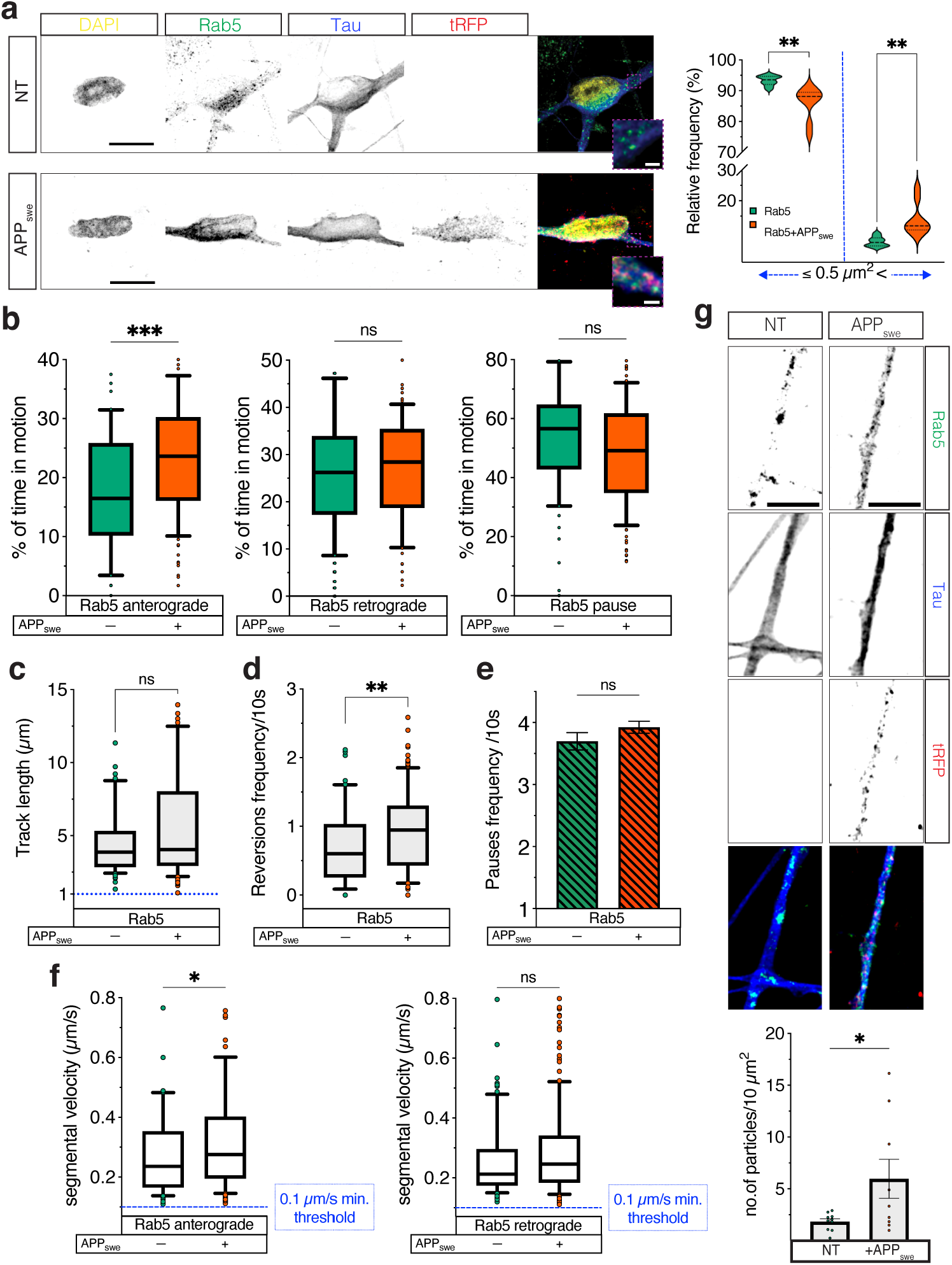
Swedish mutation perturbs axonal transport of early endosomes **a.** Representative images of NT and APP_swe_ transduced cultures stained with DAPI (blue), Rab5 (green), tRFP (red) and Tau (blue) showing different areas of Rab5+ puncta; violin plots showing puncta that are smaller or larger than 0.5 µm^2^; n≥9 cells from 3 biological replicates. **b.** Real-time movement of Rab5+ particles in NT and APP_swe_ transduced cultures moving in anterograde or retrograde direction or pausing; n≥70 particles from 3 biological replicates. **c.** Track lengths of Rab5+ particles in NT and APP_swe_ transduced cultures; n≥70 particles from 3 biological replicates. **d.** Reversions frequencies of Rab5+ particles in NT and APP_swe_ transduced cultures; n≥70 particles from 3 biological replicates. **e.** Pauses frequencies for Rab5+ particles in NT and APP_swe_ transduced cultures; n≥70 particles from 3 biological replicates. **f.** Segmental velocities of anterograde and retrograde movement of Rab5+ particles in NT and APP_swe_ transduced cultures; n>70 segments from 3 biological replicates. **g.** Representative images of Rab5+ particles in NT and APP_swe_ transduced cultures stained against Rab5 (green), tRFP (red) and Tau (blue) (scale bars= 5µm); densities of Rab5 particles in distal neurites; n≥9 cells from 3 biological replicates 0-100 percentile’s violin plot **(a)** or 10-90 percentile’s box-and-whiskers **(b, c, d, and f)**. Mean±s.e.m. **(e and g)** (*p<0.05, **p<0.01, ***p<0.001). Statistical comparisons were performed using mixed-effect analysis with Šídák’s multiple comparisons test **(a)**, Mann-Whitney test **(b, c, d, and f)** and unpaired *t*-test **(e and g)**.

We next examined the effects of the FAD Swedish mutation on the axonal transport of Rab5+ endosomes. Neurons were co-transduced either with APP_wt_/Rab5-RFP or with APP_swe_/Rab5-EGFP and axonal Rab5 transport examined. There were no differences in the number of Rab5-RFP or Rab5-EGFP tracks in any of the transduction combinations (Fig. S4b, S5a). Comparison of neurons transduced with Rab5-RFP or with Rab5-RFP/APP_wt_ also showed no differences in the axonal Rab5-RFP transport in any of the examined parameters performing either net or segmental axonal transport analyses (Fig. S5b-f). In contrast, neurons transduced with Rab5-EGFP/APP_swe_ showed significantly increased anterograde motion of Rab5 particles compared with Rab5-EGFP transduced neurons with comparable Rab5 track lengths (Fig. 5b, c, S4c). These findings were accompanied by significantly increased frequency of reversions, but not of pauses frequency, in Rab5-EGFP/APP_swe_ compared with Rab5-EGFP transduced neurons (Fig. 5d, e). Segmental velocities of anterogradely, but not retrogradely, moving Rab5 particles were also significantly increased by APP_swe_ co-transduction (Fig. 5f, S4d).

Considering the Swedish mutation increased anterograde transport of Rab5+ endosomes, we last investigated whether it also increases their density in distal projections. We observed significantly increased number of Rab5+ particles in APP_swe_ *versus* NT distal projections (Fig. 5g). These experiments indicate that the Swedish mutation impairs axonal transport of not only APP, but also of other cargoes.

## DISCUSSION

Accumulating evidence suggests that impairments in axonal transport by FAD mutations of APP contribute to the pathogenesis of AD. Animal models carrying different FAD mutations showed decreased levels of APP at the proximal stamp of ligated sciatic nerves and reduced Mn^2+^ and radiolabelled NGF transport in the hippocamposeptal pathway^25,26,31,32^. These observations indicate a role of FAD mutations in axonal transport, however, lack direct comparison between wildtype and FAD mutant APP overexpression. Consistent with these findings, cell culture experiments carrying different FAD mutations, expressing βCTFs or blocking β- and γ-cleavage sites of APP by genetic or pharmacological means all revealed perturbed axonal transport^27,33^.

We here build from these studies and characterize the axonal behaviour of APP transport elicited by the Swedish mutation, which acquires typical features of bi-directional movement. This behaviour consists of reduced anterogradely transported APP particles with slower velocities, shorter track lengths and frequent pauses, concomitant increased reversals, and time in retrograde motion. Such axonal transport behaviour is reminiscent of perturbed processivity of cargo motor assemblies^34,35^ and prompted us to investigate the interactions between APP and molecular motors. We found that the Swedish mutation enhances recruitment of DCTN1 to the APP motor assemblies. Considering DCTN1 is fundamental for processive motility of the retrograde motor dynein^3^, our finding suggests that by recruiting more DCTN1 to the APP motor assemblies, the Swedish mutation activates retrograde machinery, perturbing the physiologically predominant anterograde transport of APP. Moreover, changes in transport behaviour of Rab5 positive early endosomes indicate that the Swedish mutation perturbs axonal transport at different levels^36^. The ultimate impact of the Swedish mutation is therefore a switch in the directionality of axonal transport with APP accumulating in the cell bodies and early endosomes in distal projections. Whether accumulation of Rab5+ endosomes in distal projections, which prevents their natural maturation within the endosomal-lysosomal system, plays a role in their enlargement and malfunction in AD awaits further experimentation.

Our findings fuel the hypothesis that impairments in axonal transport underlie the axonal pathology and pathogenesis of AD^1,2,37,38^. Genetic manipulation of APP^24^, its proteolytic machinery^22,25,39^ and of all other major proteins linked to AD including tau^40,41^ and ApoE^42^ in flies and mice produces invariably axonal pathology reminiscent of the one described in molecular motor deficiencies^43^. These previous studies, together with our findings, suggest that perturbed processivity of molecular motors underlies axonal transport defects and ultimately translates into axonal pathology in AD. This hypothesis is further strengthened by the recently observed KLC1 abnormalities in the brains of patients with sporadic AD^44-46^, which suggest that besides FAD in cargos, molecular motors also play a role in axonal transport impairments. The discovery of extensively compromised axonal transport pathways is directly relevant to understand not only the pathogenesis of AD, but also of other neurodegenerative disorders.

## MATERIALS AND METHODS

### Human neuronal differentiation

Human Neural Stem Cells (hNSCs) derived from the NIH approved H9 (WA09) human embryonic stem cell line were purchased from Merck (Germany). The hNSCs were plated on matrigel-coated 100 mm petri dishes and maintained in culture with NSCs expansion media (KO DMEM/F12, 2% StemPro Neural Supplement, 1% Glutamax, 20 ng/ml β-FGF, 20 ng/ml EGF), which was exchanged every second day (DIV0-3). Upon reaching confluency, the cells were grown in the neural progenitors differentiation media (DMEM/F12, 1% B27, 0.5% N2, 1% Glutamax), which was exchanged every other day until DIV9. Neural progenitors were then detached and centrifuged at 300xg for 5’ at RT and pellets resuspended in proper Neuronal Optimized Media complete (NOMc) (DMEM-F12, 2% B27, 1% N2, 1 µg/ml laminin, 100 nM cAMP, 200 ng/ml ascorbic acid, 10 ng/ml BDNF, 10 ng/ml GDNF, 10 ng/ml IGF). Cells were counted and then seeded at different densities depending on the experimental needs. Neurons were differentiated by changing the NOMc every 6 days up to DIV40.

### Human SHSY-5Y differentiation into neuron-like population

SH-SY5Y (ATCC, USA) cells can be differentiated from a neuroblastoma-like state into a human neuronal-like cell culture. Indeed, for our experiments, we decided to differentiate the cells by applying an adaptation of a well established differentiation protocol (Shipley, M. et al., 2016.).

SH-SY5Y were seeded into 60 mm petri dishes previously coated for 15 min with 0.1 % Gelatin/PBS. Cells were maintained in DMEM complete (DMEM-high glucose, 10% Fetal Bovine Serum, 1% P/S) up to 80% confluency (DIV 0-2). Afterwards, cells were passed into new 60 mm dishes and culture medium switched to serum-free Optimem containing 1% B27, 10 ng/ml BDNF, 10 ng/ml cAMP, 10µM RA, 1% P/S, 1% Glutamax, and changed every second day up to DIV 20, when the cells were used for experiments.

### Microfluidic chambers

The microfluidic chambers (Xona Microfluidics LLC, CA, USA) were cleaned with 100% ethanol and glass coverslips coated overnight at 37ºC with a poly-ornithine solution (0.1 mg/ml in PBS). The day after, coverslips and microfluidic chambers were bonded, and the reservoirs filled with PBS to avoid bubble formation. After an hour, PBS was removed and matrigel added in all the reservoirs to fully coat both neuronal and axonal compartments. Chambers were maintained in the incubator at 37°C in 5% CO_2_ for at least 1h. Immediately before seeding, matrigel was removed and NPCs seeded in the top well of the neuronal compartment (300000 cells/50 µl in NOMc). Chambers were then placed into the incubator for 30 min, after which the wells were topped up with NOMc. Cultures were maintained by changing media every 6 days and equilibrated every 3 days to compensate for evaporation. DIV40 axons crossed completely the axonal compartment.

### Lentiviral vectors

LV-APP_wt__GFP, LV-APP_swe__tRFP and LV-GFP_Rab5 were designed to be expressed under human synapsin 1 promoter. Cloning and packaging were performed by Flash Therapeutics (France) and Vector builder (VectorBuilder Inc., IL, USA) for APP_wt_ and APP_swe_, and Rab5, respectively. The CellLight™ Early Endosomes-RFP, BacMam 2.0 expressing Rab5 was used according to manufacturer’s instructions (Thermo Fisher).

### Transduction

Human Neurons or SHSY-5Y cells were transduced with lentiviral particles at DIV17 and DIV9, respectively. Particles were retrieved from −80ºC and slowly thawed on ice for 20 min. Afterwards, specific volumes of LVs were resuspended in NOMc media according to the validated M.O.I. and TU/ml provided by the manufacturer. The transduction was performed by replacing media with either NOMc or Optimem-containing lentiviral particles for approximately 24h. Solutions containing lentiviral particles were then replaced by fresh media. Transduction levels were checked every 48 h until the start of the experiments.

### Live imaging and tracking

Movies of axonal transport of APP_wt__GFP, APP_swe__tRFP, GFP_Rab5 and RFP_Rab5 were acquired and analyzed using the same protocol. Movies were acquired at 2fps using a confocal microscope equipped with a live module (Zeiss Confocal LSM780, Zeiss Live LSM7) and an immersion oil objective 63x/1.4 NA Plan Apochromat. Time-lapse movies were processed with ImageJ prior to the analysis in Imaris (version 9.2, Oxford Instruments). Particles were tracked with the semi-automated spot tracking algorithm and visualized during the whole period of the movie.

For the analysis, we were first asked to choose which algorithm would better define the behaviour of our cargoes. Considering that studied cargos all exhibit an almost continuous movement, we applied the Autoregressive Motion algorithm. The algorithm required the input of the following parameters: XY estimated diameter, max distance, and max gap size. Diameter was based on an average empirical value for each specific cargo analyzed. For either max distance or max gap size, both spatial and temporal resolutions of the acquired movies were taken into account. Among all the statistical values obtained, the most significant one was the spatial displacement (ΔD_x_ (t_1_, t_0_) = P_x_(t_1_) – P_x_(t_2_)) of the particles in each frame and the track duration (td=total time during which a particle moves), which were both exported for further computation of axonal transport parameters. For transport dynamics analysis, we first divided the tracks into stationary or moving. All the tracks moving at < 10s were excluded. In the net axonal transport analysis, tracks with average velocities <0.10 µm/s were defined as stationary. All the other tracks were classified as moving in anterograde or retrograde direction if the average velocities were > 0.01 µm/s or < - 0.01 µm/s, respectively. In the segmental axonal transport analysis, we used the Δ_X_ displacement between frames to compute instantaneous changes in movement: pauses frequency, reversions, segmental velocities and real-time movement. Finally, since the software allows calculating the distance along each vector (x and y) of the particle’s movement, the total distances run by the particles were shown as track lengths.

### Immunocytochemistry

Neurons differentiated either in microfluidic or ibidi chambers (6 channels or 8 multi well- 8mw) were fixed in 4% paraformaldehyde (PFA)/ 4% Sucrose for 1h or 40 min, respectively. Incubation with 0.1M Glycine for 5 min was used to quench the fixation. After that, cells were permeabilized with 0.1% Tryton X-100 for 10 min. Samples were blocked with 5% BSA for 30 min and then incubated overnight at 4ºC with primary antibodies in 3% BSA. The second day, the cells were incubated with secondary antibodies in 3% BSA for 2h. Finally, cells were stained with DAPI for 3 min, then washed with ddH_2_O and mounted either with 50% Glycerol in 0.01% Na azide/PBS (ibidi 6 channels) or with Mowiol (ibidi 8mw and coverslips). Samples were dried and then stored at 4ºC.

### Confocal imaging

Fixed cells were examined either with an inverted Zeiss LSM 780 confocal microscope (Zeiss, Germany) or with a Leica DM 6000B (Leica Microsystems, Germany) using an oil immersion objective (63X/1.4 NA plan Apochromat). Z-stacks were acquired for the MAP2 and pNFH imaging analyses in both ibidi and microfluidic chambers. For APP_wt_ *versus* APP_swe_ localization, z-stacks in tile-scan mode were acquired to image APP distribution in whole neurons. To define Rab5 particle sizes, a Lightning module (Leica microsystems) was used to deconvolve z-stacks and improve particle resolution for an unbiased analysis of the puncta size.

### Identification of the axonal nature of the neurites

Ibidi µ-Slides VI ^0.4^ were used to perform part of the transport experiments for both APP_wt__GFP and APP_swe__tRFP. Differently from the microfluidic chambers, the ibidi device lacks a physical division between the neuronal and axonal compartments. For this reason, prior to seeding the neurons, the ibidi were marked with an arbitrary reference point to set an X; Y (0;0), this was then matched with the 0;0 of the confocal stage prior to time-lapse acquisition. While acquiring movies, each selected projection was marked into a position list and saved. Post-live imaging cells were used for immunocytochemistry, following the protocol described above, and positions retrieved to identify exclusively pNFH(+) neurites, which were included into the axonal transport analysis.

### Image analysis

Images acquired from immunocytochemistry were analyzed either with ImageJ or Imaris.

i. <u>MAP2/pNFH characterization in human neurons:</u> from 40 z-stacks acquired per each different biological replicate (n=3), 20 images were randomly selected for further ratio analysis. Among the 20 pictures, 5 randomly selected neurites, either MAP2-(+) or pNFH- (+) were selected in each channel and intensities measured to compare the ratios among all the samples.
ii. <u>pNFH-GFP/tRFP post-live imaging:</u> neurites were traced in ImageJ following GFP or tRFP intensities for APP_wt__GFP and APP_swe__tRFP, respectively. Intensity profiles of APP were matched with those of pNFH to either include or exclude the neurite from further axonal transport analysis.
iii. <u>APP localization:</u> ROIs were traced for both cell bodies and neurites to measure either APP or Tau intensities. Tau was used as a reference marker to study both APP_wt_ and APP_swe_. Afterwards, ratios of APP in soma/axon were measured and normalized for Tau intensity. With the same approach, Tau levels in both APP_wt_ and APP_swe_ were compared by quantifying the soma/axon ratios.
iv. <u>Rab5 size and densities:</u> Rab5 puncta areas were measured in deconvolved z-stacks. Tau was used as a mask. When transduced with APP_swe_, only those cells that were overexpressing mutant APP were used to measure Rab5 sizes. Masked images were analyzed in Rab5 channel by the Analyze particles tool from ImageJ to determine the area of the puncta in the whole neuron, and their densities in distal projections.

### Protein extraction

For Immunoprecipitations (IPs), proteins from differentiated SHSY-5Y cells, Non-Transduced (NT) or transduced with either APP_wt__GFP or APP_swe__tRFP, were collected with the IP lysis buffer (1% Nonidet – P40, 25 mM Tris buffer pH 7.4, 150 mM NaCl, 1mM EDTA, 5% Glycerol). Cells were incubated for 30 min on ice and cell membranes disrupted using an insulin syringe. Supernatants were collected into new tubes following centrifugation for 20 min at 20000xg 4ºC. Total lysates were quantified using the BCA assay (Pierce™ BCA Protein Assay Kit, Thermo Fisher).

### Immunoprecipitations

Target proteins were immunoprecipitated either with GFP- or RFP-trap Magnetic Agarose beads (Chromotek, Proteintech) or with Dynabeads Protein G (Thermo Fisher), which both allow magnetic separation of target proteins. GFP or RFP-trap were used to IP APP_wt__GFP and APP_swe__tRFP, respectively. Prior to the addition of proteins, beads were equilibrated with 500 µl of Dilution buffer (10 mM Tris/Cl pH 7.5, 150 mM NaCl, 0.5 mM EDTA, 0.018% Na azide) as instructed by the manufacturer’s protocol. 1 mg of proteins was used per each IP. Beads were used at 50 µl/mg of total lysate for both GFP and RFP trap and samples incubated for 2h at 4ºC rotating end-over-end. Tubes were spun at 1000xg for 1 min and then placed into DynaMag-2 (Thermo Fisher) for separation of flow-through fractions. The beads were washed twice for 2 min with 500µl/wash buffer (150 mM NaCl, 50 mM Tris/Cl pH 7.5). Protein complexes were eluted using 1X LDS sample buffer, 1X NuPage DTT reducing agent (Thermo Fisher) and heated at max 80ºC for 15 min. Eluates were last saved into new tubes for SDS-PAGE and WB blot analysis.

For IP of DCTN1, 50µl of Dynabeads Protein G were used per sample. The beads were freed from storing solution by magnetic separation and then the antibody conjugation performed by adding 200µl of the Ab Binding Buffer (Thermo Fisher), and 5 µg of DCTN1 antibody. The mix was incubated for 1h rotating end-over-end at 4ºC. Ab Binding Buffer was then discarded and 1mg of total lysate added to the beads. Incubation, washes, and elution were performed as above (washes were made in this case by using just 200 µl of Wash Buffer).

### SDS-PAGE and Western blotting

Quantified protein samples were loaded into Bolt Bis-Tris Plus 4-12 % precast gels (Novex, Thermo Fisher) and SDS-PAGE performed first for 10 min at 50V and then at 110V for 1h 30 min. Proteins were then transferred onto PVDF membranes (Thermo FIsher) using a Mini blot module (Invitrogen) for 1h 50 min at 30V. Membranes were washed 3 × 5 min with 20mM Tris-Buffer (TBS) and blocked in 5% non-fat dry milk (NFDM) in 20mM TBS/0.2% Tween-20 (TBS-T) for 1h at RT. After 3 washes in TBS, the membranes were probed overnight at 4ºC with primary antibodies listed below in 1% BSA in TBS-T. The second day, HRP-conjugated antibodies were prepared in 1% NFDM TBS-T and incubated for 2h at 4ºC. Last, the membranes were washed 3 × 5 min with TBS-T and proteins detected using Chemiluminescent Substrate (SuperSignal™ West Pico PLUS, Thermo Scientific) by acquiring images with Chemidoc (BioRad).

### Antibodies

The following antibodies were used for immunocytochemistry: MAP2 (1:3000, Abcam 221693), pNFH (mouse 1:2000, BioLegend 801601). TurboRFP (1:1000, Evrogen AB233), GFP (1:1000, abcam 5450), Tau (1:400, abcam 62639) and Rab5 (mouse 1:1000, Cell Signaling 46449). For IP and blots we used APP (1:1000, Abcam ab126732), KLC1 (1:1000, Abcam 174273), DCTN1 (mouse 1:200 Santa Cruz 135890, rabbit 1:1000 or 1:50 Cell signaling 69399) and β III-tubulin (1:6000 Biolegend 802001). Secondary antibodies used for immunocytochemistry were donkey anti-mouse/rabbit/goat/sheep conjugated to Alexa Fluor 488/546/555/647 (Invitrogen 1:500), respectively. For immunoblotting, antibodies conjugated with HRP were used: anti-Rabbit 1:2000 (Cell signaling 7074), anti-Mouse 1:2000 (Cell signaling 7076), anti-Goat 1:1000 (Santa Cruz 2354), and Clean-Blot™ IP Detection Reagent 1:200 (Thermo Scientific 21230).

### Statistical analysis

All the analyses were performed by using GraphPad Prism 9.4.0. Statistical tests are described in detail in figure legends.

## ACKNOWLEDGEMENTS

We thank members of the Stokin Lab for support and feedback. Among all we thank in particular Dr. V.M. Pozo Devoto for data analyses and N. Dragišić for assisting with cell cultures.

## AUTHORS CONTRIBUTIONS

Conceptualization, M.F. and G.B.S.; Methodology, M.F. and G.B.S.; Investigation, M.F.; Formal Analysis, M.F.; Data Curation, M.F. and G.B.S.; Writing, M.F. and G.B.S.; Supervision, G.B.S.; Project Administration, G.B.S.; Funding Acquisition, G.B.S.

