## Supplementary Figures and legends for "Familial Alzheimer’s disease mutation undermines axonal transport by enhancing dynactin recruitment to the APP motor assemblies"

#### Fig.S1 – FAD mutations of APP impair axonal transport

**a.** Number of tracks detected in APPwt, lon or swe projections; n=8 distal projections.

**b.** Proportions of $APP_{wt}$, $APP_{swe}$ and $APP_{lon}$ in anterograde or retrograde movement and stationary; n=8 distal projections.

**c.** Average velocities of $APP_{wt}$, $APP_{swe}$ and $APP_{lon}$ in anterograde or retrograde movement; n=8 distal projections.

**d.** Representative image of the microfluidic chamber and of the immunostaining of human neurons labelled with pNFH and MAP2 (scale bar=300 µm).

**e.** Proportions of $APP_{wt}$ or $APP_{swe}$ in anterograde or retrograde movement and stationary measured from the axons grown in microfluidic chambers; n≥4 axons from 3 biological replicates.

Data are mean±s.e.m. (a, b, and e) or 10-90 percentile's box-and-whiskers (c). Statistical comparisons were performed using one-way ANOVA with Tukey's multiple comparisons test (a) or Dunnet's multiple comparison's test (b), Kruskal-Wallis test (c) and mixed-effect analysis with Šídák's multiple comparisons test (*p<0.05, ***p<0.001).

#### Fig.S2 – Segmental analysis of the APP axonal transport

**a.** Total tracks distances of axonal $APP_{wt}$ and $APP_{swe}$ n>150 tracks from 3 biological replicates.

**b.** Reversions frequencies of axonal $APP_{wt}$ and $APP_{swe}$ examined separately for anterograde and retrograde tracks; n>45 tracks from 3 biological replicates.

**c.** Pauses frequencies of axonal $APP_{wt}$ and $APP_{swe}$; n>150 particles from 3 biological replicates.

**d.** Mean segmental velocities of axonal $APP_{wt}$ and $APP_{swe}$ examined separately for anterograde and retrograde segments; n>400 segments form 3 biological replicates.

Data are 10-90 percentile's box-and-whiskers **(a-c)** or mean±s.e.m. **(d)**. Statistical comparisons were performed using Mann-Whitney test **(a-c)** and unpaired *t*-test (**d**, *p<0.05, ***p<0.01).

**Fig.S3 – DCTN1 recruitment to APP motor assemblies is caused by Swedish mutation**

**a.** Biochemical analyses with anti-GFP and anti-tRFP IPs reveal that both DCTN1 and KLC1 co-immunoprecipitate with either $APP_{wt}$ or $APP_{swe}$ complexes.

**b.** Membranes were loaded with Total Lysate (TL) or IP-GFP from NT or $APP_{wt}$ transduced cells and probed with APP (lower blot) or GFP (upper blot), respectively.

**c.** Membranes were loaded with Total Lysate (TL) or IP-tRFP from NT or $APP_{swe}$ transduced cells and probed with APP (lower blot) or tRFP (upper blot), respectively.

**d.** Biochemical analyses show co-IP between DCTN1 and both $APP_{wt}$ and $APP_{swe}$.

**Fig.S4 – Segmental axonal transport parameters of Rab5 that are not changed in $APP_{swe}$ cultures**

**a.** Areas of Rab5 puncta in NT and $APP_{swe}$ cultures; n≥9 neurons; unpaired t-test.

**b.** Number of axonal Rab5 tracks in ± $APP_{swe}$ cultures; n>5 axons from 3 biological replicates.

**c.** Proportions of moving or stationary Rab5 particles in ± $APP_{swe}$ cultures; n>5 axons from 3 biological replicates.

**d.** Frequency distributions of anterograde and retrograde segmental velocities depicted using cubic spline fit of Rab5 particles in ± $APP_{swe}$ cultures; n>70 from 3 biological replicates.

Data are mean±s.e.m. **(a-c)** or frequency distributions **(d)**. Statistical comparisons were computed using unpaired *t*-test (a-c) and Mann-Whitney test (**d**, for p-values see **fig.5f**) (*p<0.05).

**Fig.S5 – Axonal transport of Rab5 particles is not perturbed by $APP_{wt}$**

**a.** Number of axonal Rab5 tracks in ± $APP_{wt}$ cells; n≥4 axons.

**b.** Proportions of Rab5 particles in ± $APP_{wt}$ cells, examined separately for moving and stationary; n≥4 axons.

**c.** Proportions of real-time moving and pausing Rab5 particles in ± $APP_{wt}$ cells; means±s.e.m; n>50 particles from 3 biological replicates.

**d.** Track lengths of Rab5 particles in ± $APP_{wt}$ cells; n>50 particles from 3 biological replicates.

**e.** Reversions frequencies of Rab5 particles in all tracks of ± $APP_{wt}$ cells; n>50 particles from 3 biological replicates.

**f.** Pauses frequencies of Rab5 particles with and without $APP_{wt}$ cells; n≥4 axons.

Data are mean±s.e.m. **(a, b, and f)** or 10-90 percentile's box-and-whiskers **(c, d and e)**. Statistical comparisons were performed using unpaired *t*-test **(a, b, and f)** and Mann-Whitney test **(c, d and f)**.

Figure S1 Feole and Stokin

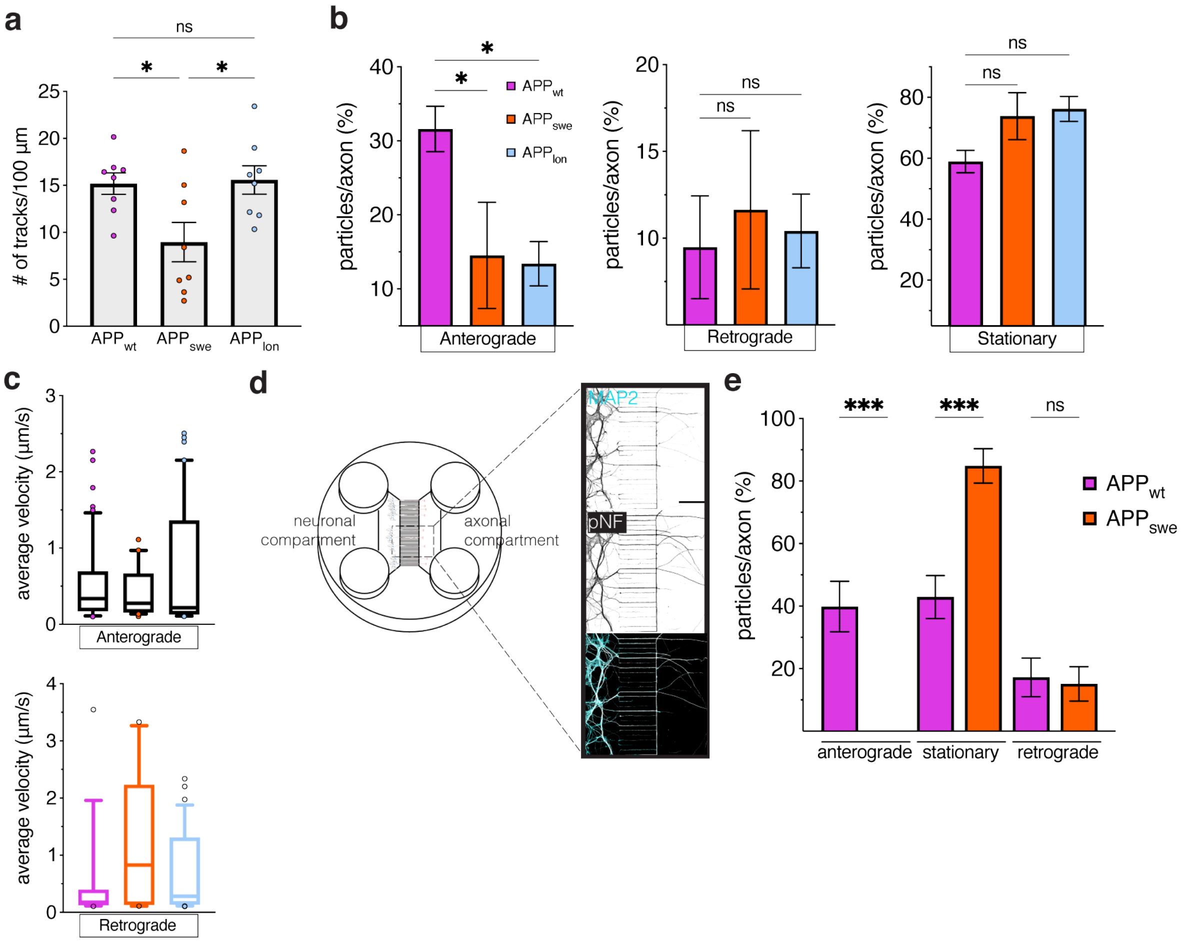

Figure S2 Feole and Stokin

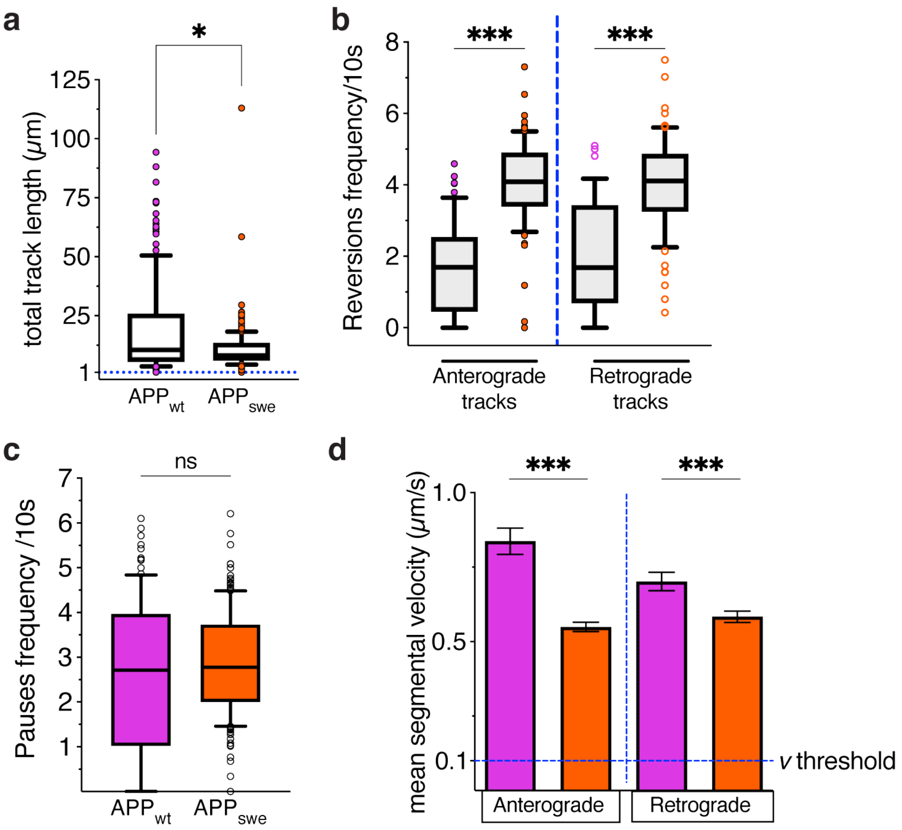

### Figure S3 Feole and Stokin

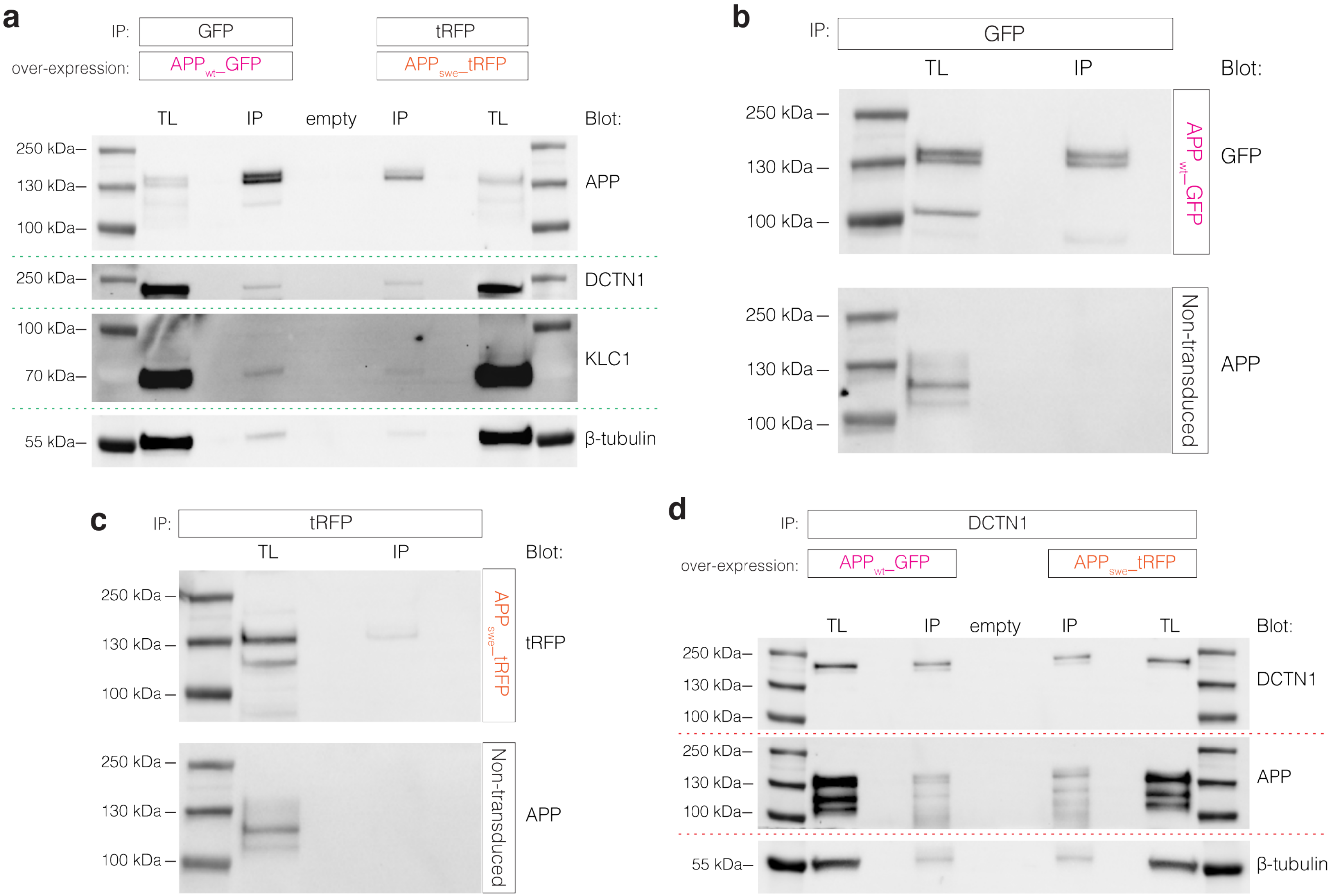

Figure S4 Feole and Stokin

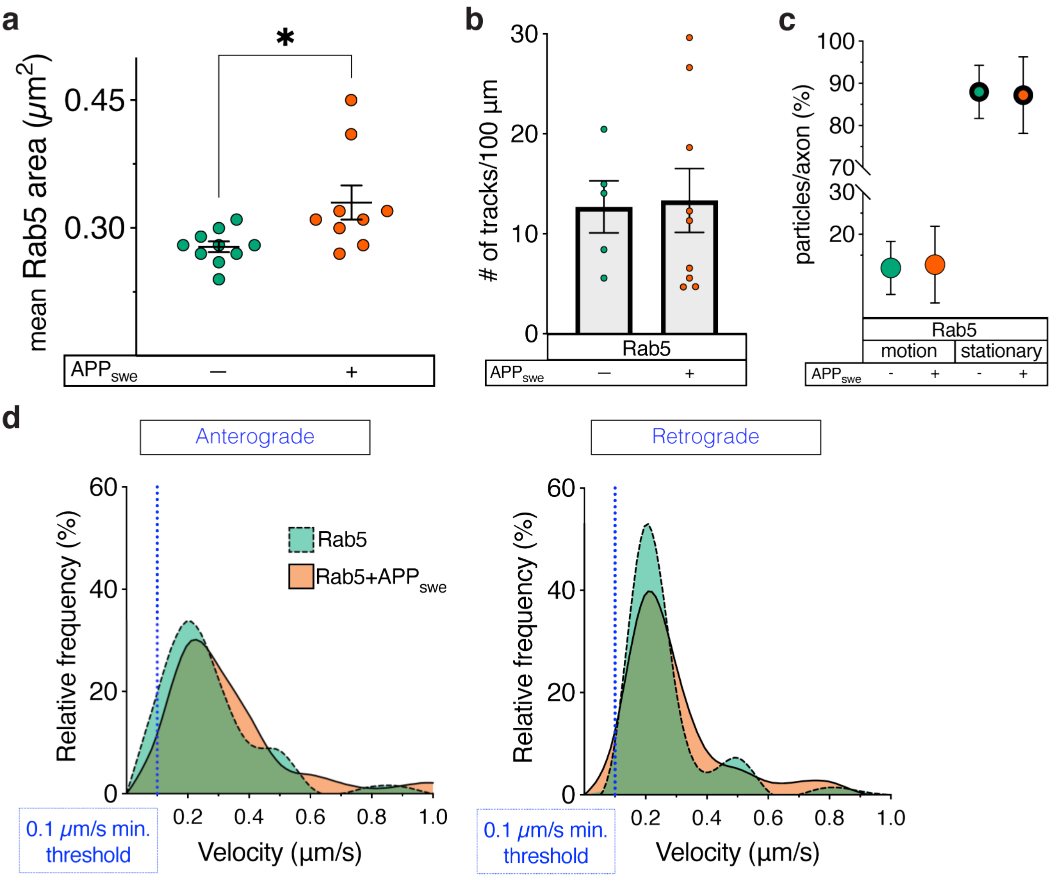

Figure S5 Feole and Stokin

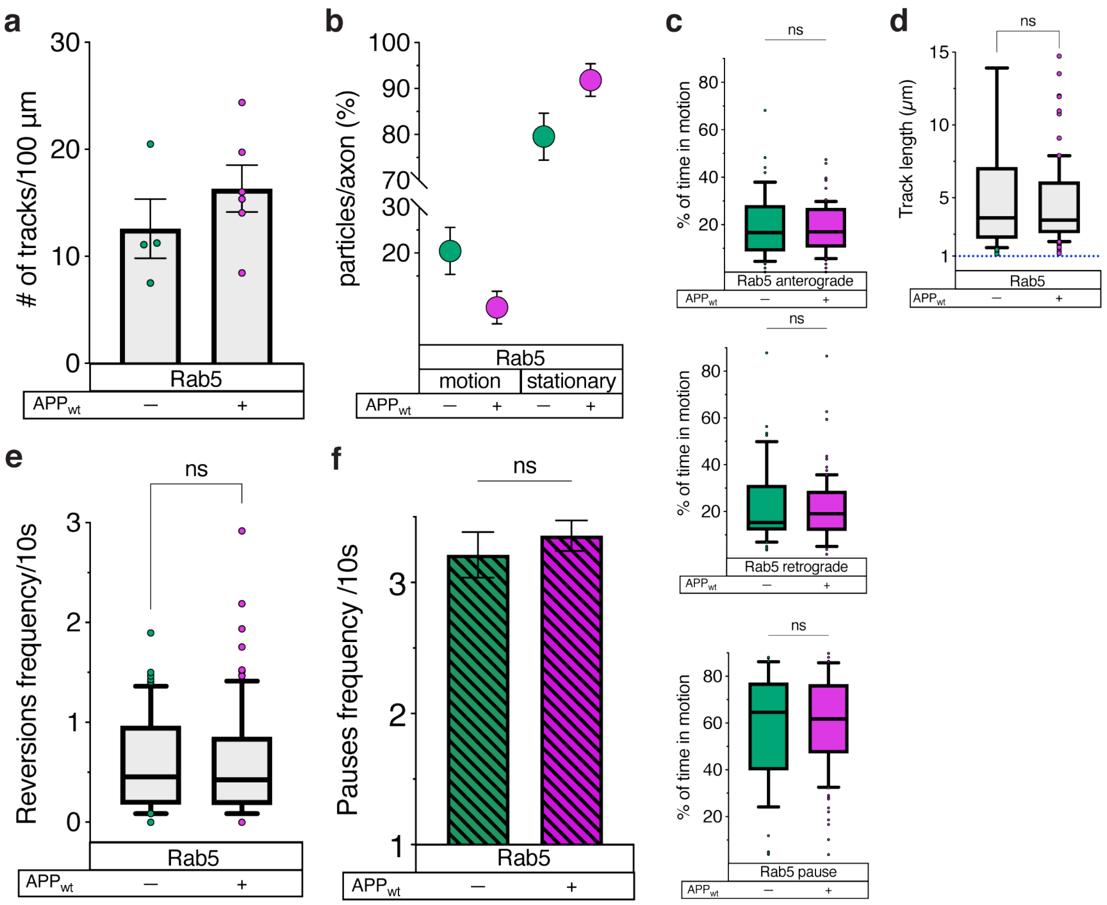
